# Neurocognitive processing efficiency for non-alarm rather than alarm signaling in human scream calls

**DOI:** 10.1101/2020.05.04.076463

**Authors:** Sascha Frühholz, Joris Dietziker, Matthias Staib, Wiebke Trost

## Abstract

Across many species, scream calls signal the affective significance of events to other agents. Scream calls were often supposed to be of generic alarming and fearful nature to signal potential threats including their instantaneous, involuntary, and accurate recognition by perceivers. However, scream calls are more diverse in their affective signaling nature than being limited to fearfully alarm a threat, and thus the broader sociobiological relevance of various scream types is unclear. Here we used four different psychoacoustic, perceptual decision-making, and neuroimaging experiments in humans to demonstrate, first, the existence of at least six generic and psycho-acoustically distinctive types of scream calls of both an alarming and a non-alarming nature, rather than being limited to only screams caused by fear or aggression. Second, based on perceptual and processing sensitivity measures for decision-making during scream recognition, we found that alarming screams (with some exceptions) were overall discriminated the worst, were responded to the slowest and were associated with the lower perceptual sensitivity for their recognition compared with non-alarm screams. Third, the neural processing of alarm compared with non-alarm screams during an implicit processing task elicited only minimal neural signal and connectivity in perceivers, contrary to the frequent assumption of a threat processing bias of the primate neural system. These findings show that scream calls are more diverse in their signaling and communicative nature in humans and that especially non-alarming screams, and positive screams in particular, seem to have higher efficiency in the cognitive, neural, and communicative processing in humans.

## INTRODUCTION

Vocal affect bursts, such as crying, grunting, or laughing, are a major part of the sociobiological communication across many mammalian species, especially primates. A specific type of affect burst is a scream call. Screams are relatively short, loud and intense, high-pitched, tremulous, and rough voice calls [1–3]. They have a far-reaching impact [1,4] and seem to be immediately recognized by, adaptively and rapidly responded to, and hardly ignored by perceivers [5,6]. Screams are thus assumed to be an effective mode of communicating affect signals that are of high relevance in any sociobiological interaction [1]. Scream calls have a long evolutionary trajectory across many species up to human primates. In nonhuman primates [7–9] and other mammalian species [10], scream-like calls are frequently used as some specific type of alarm signal exclusively in negative contexts, such as in in-group social conflicts between rank different animals. Screams by lower ranking animals help to recruit support from allies [11,12], while higher ranking animals scream to intimidate the lower ranking [13]. Furthermore, “SOS screams” signal the presence of environmental threat (e.g. predators) [14,15]. Scream-like voice calls thus aim to trigger certain behavior in potential listeners, similar to other types of alarm calls that are commonly expressed in high-arousal states of fear [16] or aggressiveness [17].

Accordingly, human screams are assumed to share this acoustical and motivational nature of screams with other species, highlighting the alarming quality of such affect bursts to signal danger and to scare potential predators [1,4]. As being of alarming nature, screams thus demand urgent responses in listeners, implying a fast recognition and an accurate perceptual categorization [5,18] as well as neural efficiency in processing [1]. This fast and efficient processing of the alarming quality of screams might be based on a certain acoustic scream feature, since a previous report [1] suggested that a specific frequency range of the temporal modulation rate was related to the perceptual feature of “roughness” (i.e. a harsh, thrilling, croaky sound) in the broad 30-150Hz range that contributes to the alarming quality of fear screams [19]. Other studies confirmed that roughness, amongst other important acoustic and perpetual features, is a defining perceptual feature of screams that enable listeners to classify vocalizations as screams [3].

Although previous studies thus provided a detailed functional and neural description of scream call recognition, they were limited from three perspectives. The first limitation concerns the variety of primate screams, especially human screams. These studies largely focused on alarming fear screams [1,5,10] and sometimes on scream-like and alarming aggressive roars [17]. However, humans scream not only when they are fearful and aggressive, but also when they experience other affective states, similar to a variety of emotions that are more commonly expressed in less intense nonverbal vocal emotions [20], such as sadness, joy, and sensual pleasure. Second, although some previous studies seemed to have investigated a broader variety of scream-like vocalizations, these vocalizations seemed intermingled with other non-scream-like vocalizations [21] or partly collected from mixed sources [3,21]. The third limitation concerns the neural mechanisms and dynamics of scream perception, as only a limited description of neural mechanisms for scream perception is as yet available [1]. To overcome these limitations, we herein aimed to provide a broader description of human screams on three largely interdependent levels that are relevant for any kind of vocal communication. These three levels are represented, first, by the acoustic properties and acoustic distinctiveness of scream calls as produced by a sender (experiment 1), second, by the perceptual and categorization dynamics of these scream calls in listeners (experiment 2-3), and, third, by the neural mechanisms of scream recognition in the central nervous system of listeners (experiment 4).

Besides these aforementioned limitations, the experiments described here were furthermore motivated by the assumption that a broader taxonomy of generic and distinct human screams potentially needs to distinguish at least (a) screams when experiencing intense pleasure, (b) desperate scream-like cries during sadness, (c) screams of joy and elation, (d) screams of pain, (e) screams of anger and rage, and (f) fearful screams (Fig. 1; Fig. S1). This broader taxonomy of screams is proposed based on a survey of many daily natural, social, and cultural manifestations of human screams, and the general diversity of human affective vocalizations [22] and especially of nonverbal expression of emotions [20]. While we assume that screams d-f are of an alarming nature (i.e. they call for immediate action), screams a-c are largely non-alarming (i.e. no fast response required; see also below). Furthermore, while screams of joy and pleasure are of a positive nature, the other four types of screams have a negative affective valence.

**Figure 1.**
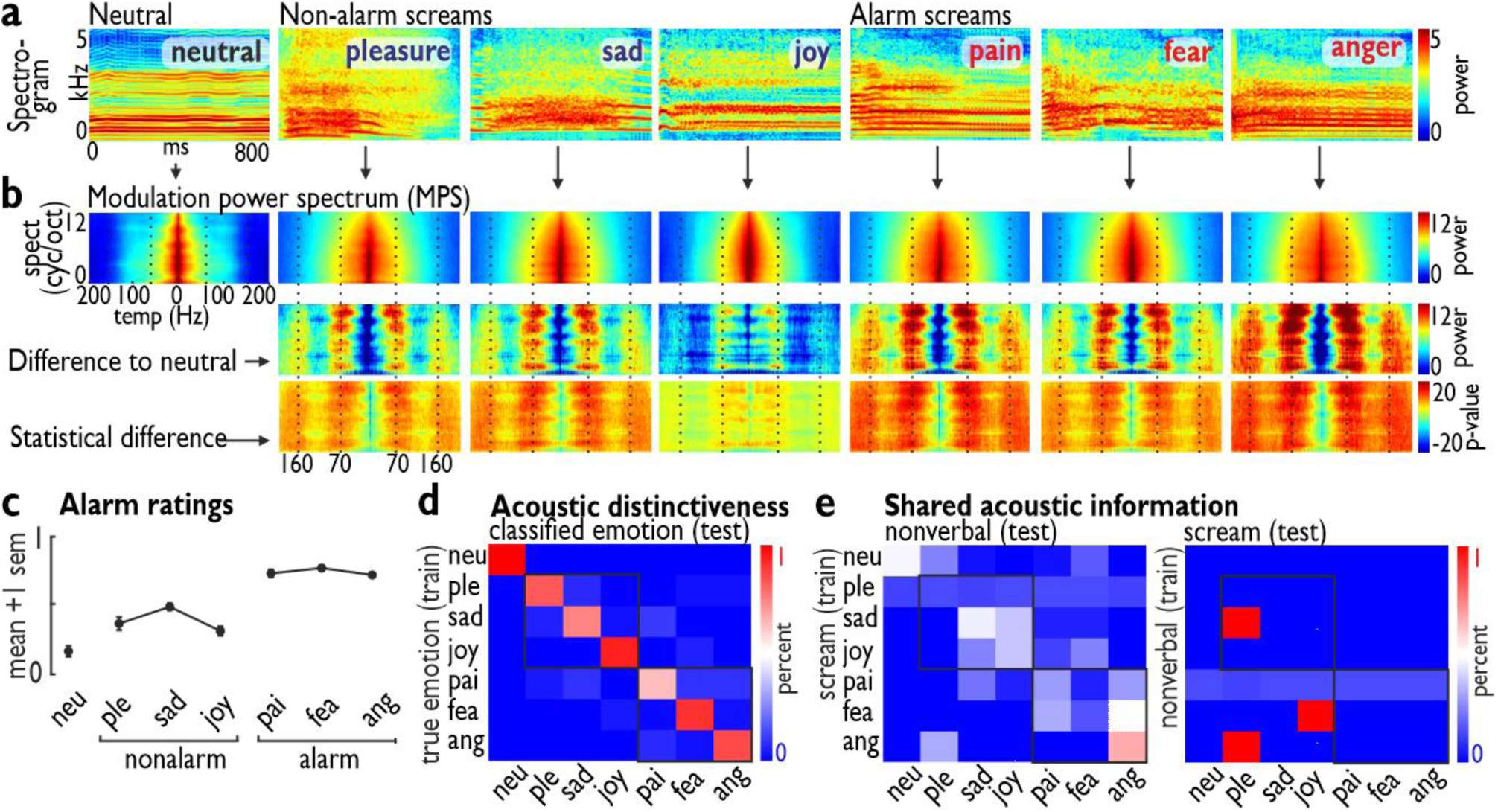
Acoustic description and alarm ratings of 6 generic types of screams. (a) Spectrograms of example stimuli for each generic scream type, including a neutral scream of an intense vocalization of the vowel /a/. (b) Average modulation power spectra (MPS) for each scream type (upper row) as the numeric (middle row) and statistical difference (lower row; n=60 per scream type; n=420 screams in total) of each generic scream from neutral screams. (c) Alarm level ratings confirmed the differential level of alarm of screams that seem to be of a low alarming nature (non-alarm screams: pleasure, sad, joy) and a high alarming nature (alarm screams: pain, fear, anger). Neu = neutral; ple = pleasure; pai = pain; fea = fear; ang = anger. (d) Using a support vector machine (SVM) (train=training data, test=test data) on the acoustic features of each scream allowed us to separate each scream type from other screams with high accuracy (e.g. low off-diagonal misclassifications). (e) To test whether screams share acoustic information with common nonverbal affect bursts, we trained the SVM either on the screams (left) or on the nonverbal affect bursts (right) and tested the classifier on the other type of affective vocalizations; in both cases, the SVM classifier failed to show superior performance (e.g. high off-diagonal misclassifications).

## RESULTS

### Psychoacoustic and affective diversity of human screams

Given the assumed communicative relevance of diverse types of scream calls, they should have a distinct and differential acoustic profile during their expression in senders as the basis for any consequent perception in listeners. Concerning this acoustic and expressive nature of screams, we accordingly performed a first experiment (experiment 1) wherein we asked human participants to vocalize a broad variety of positive and negative screams from instructions of typical situations for each type of scream. Participants (n=12) vocalized these 6 types of screams in addition to a neutral but intense vocalization of the vowel /a/. This category of this neutral vocalizations (referred to as “neutral screams” for the sake of terminological simplicity) was included as baseline and comparison condition to quantify acoustic, perceptual, and neural differences in processing the 6 generic vocal screams. The recordings of the neutral and the 6 generic scream types resulted in a total of 420 acted scream calls that are acoustically similar to natural screams [1,2]. The screaming quality of these vocalizations was verified by ear by trained senior members of the research team, since clear formal acoustic criteria for scream calls are not yet established [3]. Scream calls are also easily identified and discriminated from other nonverbal affective vocalizations [21].

To test whether all screams are always of a high alarming nature, we asked an independent sample of participants (n=23) to rate the alarming quality of each scream. Although the term “alarm” often refers to certain sociobiological contexts and events that elicit screams especially in animal settings, the scream can be regarded as the senders’ expression of the alarming significance of the context or event. Listeners than rate the alarming level of these screams as part of their decision how urgent one needs to respond. We quantified these listeners alarm rating individually for each scream and on a dimensional basis (i.e. intendent of the knowledge to which category the scream belonged to). As hypothesized, screams differed significantly in their alarming quality (1-way analysis of variance [1w-ANOVA], 7 levels; F_1,6_=113.95, p<0.001, η^2^=0.77; Greenhouse-Geisser (GG) correction applied to all p-values in case of sphericity violations based on a Mauchly’s test), such that alarm screams (pain, anger, fear) were overall significantly more alarming (1w-ANOVA, 3 levels; F_2,44_=227.63, p<0.001, η^2^=0.74) than non-alarm screams (pleasure, sad, joy) (p<0.001) and neutral screams (p<0.001) (Fig. 1c). We therefore categorized the pain, anger, and fear screams as “alarm screams” and pleasure, sad, and joy screams as “non-alarm screams”. This categorization was based on a full permutation and statistical testing through any possible categorical combination of the 6 generic screams (see methods). Categorizing all screams into neutral screams, non-alarm screams (pleasure, sad, joy), and alarm screams (pain, anger, fear) led to the overall most significant result (1w-ANOVA, 3 levels; F_1,2_=227.63, p<0.001, η^2^=0.84), while any other kind of combination was less significant (all F’s_1,2_=114.24-225.91). The chosen categorization thus maximized the difference between the scream categories along the alarming dimension the best, followed by the second-best categorization of only fear and anger as alarm screams (F_1,2_=225.91), and the third-best categorization of only fear and pain as alarm screams (F_1,2_=221.23).

Besides this perceptual alarm rating of the scream calls, we also performed an acoustic analysis of each type of scream. In a first acoustic analysis, we calculated the spectrogram of each of the alarm and non-alarm screams (Fig. 1a) and then quantified the modulation power spectrum (MPS) [23,24] of the three alarm and the three non-alarm screams (Fig. 1b). The MPS is an estimate of the spectral and temporal information in the acoustic scream signal, and was calculated similar to a previous study [1]. We found that these temporal modulation rates were focused around frequency bands centered on low level ∼60Hz (range 50-70Hz) and a high level ∼160Hz (range 140-180Hz), and this was characteristic both of alarm screams (fear, pain, anger), and most important also of some non-alarm screams (pleasure, sad). These modulation rates for the temporal evolvement of sound have been thought to be associated with the perceptual features of “roughness” [1]. Although this roughness is not an exclusive feature of screams, as many other nonverbal vocalizations can have a rough acoustic attribute [21], it is nonetheless very characteristic of screams. Specifically, the power of the modulation rate quantified for each scream in the low (ß=15.665) and high-frequency range (ß=4.903) predicted positively the alarm ratings for each scream as quantified by a linear regression analysis (F=138.240, p<0.001, R^2^=0.399).

Many types of alarm and non-alarm screams thus seem to share a rough acoustic property, but roughness is only one feature among many other important voice features. We therefore asked in a second step of the acoustic analysis whether scream types are acoustically different overall beyond this roughness feature. We therefore analyzed each scream type according to a larger and well-established set of 88 acoustic features [25], including a proper tracking of acoustic pitch feature (see methods). These 88 features are centrally related to a general acoustic voice space. We subjected these features to a machine learning approach implemented as a support vector machine (SVM). The SVM was trained (train data) on the acoustic features of a part of the screams, and then was tested (test data) to classify a new set of screams into 7 possible categories (Fig. 1d). The overall classification accuracy in this cross-validation approach was high at 79.8% (chance level 14.3%), with accuracies ranging from 65.8% (sad) to 89.7% (joy) and 100% (neutral). This confirms that each type of scream is acoustically very distinctive from each other at a high level, and also that non-alarm screams are equally distinctive as alarm screams, within and across these larger scream categories.

We finally also tested whether the 6 scream types are acoustically distinct from more basic nonverbal affect vocalizations of similar affective valence, such as laughs, cries, or anger growls [26]. In this cross-classification approach, despite training the SVM on scream calls, it was unable to classify nonverbal vocal affect (overall classification accuracy 14.3%, chance level 14.29%), and the acoustic features of nonverbal affective voices did not help with classifying screams (overall accuracy 19.2%) (Fig. 1e).

### Low perceptual sensitivity for alarming screams

Expressing scream calls by senders is the basis for their communication and for the potential detection and recognition by receivers. Having shown in experiment 1 that human screams are not limited to alarming screams signaling threat, but rather show an acoustic diversity of at least 6 different types of screams, we investigated in experiment 2 how accurately human listeners can perceptually discriminate and classify these different scream types. We therefore presented a selection of 84 of the original 420 scream calls (see methods; all selected screams had an equal base recognition rate) to another sample of human listeners (n=33) and asked them to classify the screams into 7 categories that referred to the 6 scream types and the neutral vocalizations (Fig. 2a).

**Figure 2.**
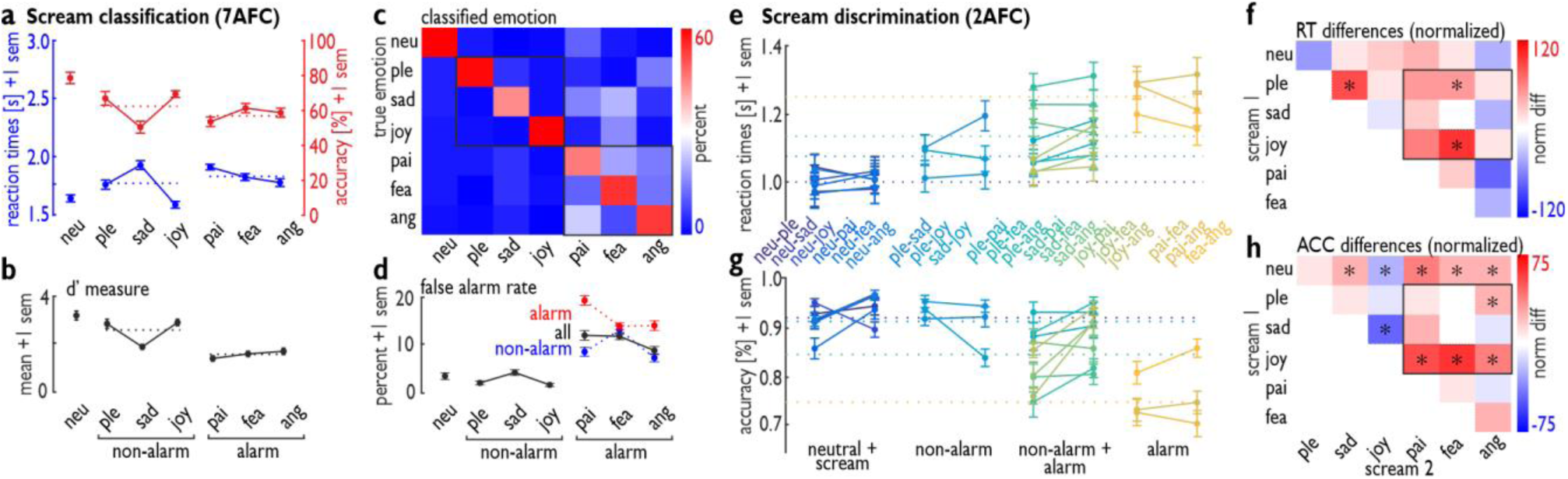
Perceptual decision-making on perceived scream calls. (a-d) Reaction times and accuracy level for (a) categorizing the 7 types of screams in a 7-alternative forced-choice (7AFC) task (top left) in experiment 2. (b) During misclassification of screams (top right, off-diagonal), participants significantly used more categories from alarm screams than from non-alarm screams (bottom right). (c) Combining the hit rate and the false alarm rate results in the d′ measure of sensitivity to detect a certain scream type (bottom left). (d) The false alarm rate is also reported separately for alarm (red) and non-alarm screams (blue). Neu = neutral; ple = pleasure; pai = pain; fea = fear; ang = anger. (e-h) Reaction time (RT) (upper panel, e) and accuracy level (ACC, lower pane, fl) for discriminating scream calls (experiment 3) from neutral screams (dark blue, left side of bar plots), for discriminating alarm screams from non-alarm screams (third column, turquoise bars), and for discriminating alarm screams from alarm screams (right side, yellow). The right side (g-h) shows the normalized RT and accuracy difference between the 21 combinations of screams. *p<0.05, based on a non-parametric permutation test.

Reaction times in correctly classified trials (1w-ANOVA, 7 levels; F_6,192_=26.228, p<0.001, η^2^=0.29) and performance accuracy rates (1w-ANOVA, 7 levels; F_6,192_=12.082, p<0.001, η^2^=0.21) significantly differed across the 7 scream types. Posthoc tests on reaction times showed that all generic screams were classified slower than neutral screams (all p’s<0.035), except for joy screams (p=0.400); all non-alarm screams had different reactions times (all p’s<0.008), while only pain and anger differed (p=0.005) in reaction times for alarm screams. Comparisons across non-alarm and alarm screams showed that joy scream were classified faster than all alarm screams (all p’s<0.001) and pleasure being classified faster than pain screams (p=0.013). Only sad screams were classified slower than anger screams (p=0.010). Postdoc tests for the performance accuracy showed that neutral screams had better a classifications rate than the negative screams (all p’s<0.001), but not compared to the positive screams (all p’s>0.103); within the non-alarm screams, sad screams differed from pleasure (p=0.034) and joy screams (p<0.001), while alarm scream did not differ in performance accuracy (all p’s>0.142). Comparisons across non-alarm and alarm screams showed that joy screams were more accurately classified than anger (p=0.004) and pain screams (p<0.001).

To better qualify these differences on a broader level, we repeated the analysis with only 3 categories on the basis of the aforementioned observation that scream calls largely differ in their alarming level. The scream calls were divided into 3 major categories, represented by the neutral screams, non-alarm (pleasure, sad, joy), and alarm screams (pain, fear, anger), and we calculated the mean reaction times and classification accuracies for each participant within these three categories. These 3 categories differed in reaction times (1w-ANOVA, 3 levels; F_2,64_=27.636, p<0.001, η^2^=0.20), with non-alarm (p<0.001) and alarm screams (p<0.001) having slower mean reaction times than those for neutral screams, and non-alarm screams being classified faster than alarm screams (p=0.005) (Fig. 2a). The latter difference however seems strongly driven by fast response to joy screams, and partly also to pleasure screams compared the other three alarm screams. These 3 categories also differed in accuracy rates (1w-ANOVA, 3 levels; F_2,64_=27.365, p<0.001, η^2^=0.28), with lower accuracy for non-alarm (p<0.001) and alarm screams (p<0.001) than for neutral screams, but with no difference in accuracy for non-alarm and alarm screams (p=0.192) (Fig. 2a).

In addition, we quantified the false alarm rate, meaning how often each scream category was selected during misclassifications (Fig. 2b,d). This analysis figures as an indicator of the decisional relevance of scream categories in subjective (mis-)classifications. The frequency of selecting alarm scream categories during misclassifications was higher (1w-ANOVA, 3 levels; F_2,64_=74.528, p<0.001, η^2^=0.43) compared with that for the neutral (p<0.001) and non-alarm categories (p<0.001) in the comparison of three major scream categories (neutral, non-alarm, alarm) as mentioned above. Thus, participants tend to perceive screams as “alarming” when they misclassify scream calls, that is, they more likely choose one of the alarming scream types during misclassification. This was true during misclassifications of both non-alarm screams (i.e. classified in one the three alarm scream types) and especially of alarm screams (i.e. classified in one of the other two alarm scream types instead of the target alarm scream type) (Fig. 2d).

By considering the false alarm rate for using a certain scream type for decisional misclassification in relation to the correctly identified scream for this category, we calculated the d′ measure as an indicator of perceptual sensitivity to a certain scream type of the 6 generic screams (Fig. 1c). All scream types differed in their perceptual sensitivity across the 7 categories (1w-ANOVA, 7 levels; F_1,6_=39.388, p<0.001, η^2^=0.48), with alarm screams revealing the lowest perceptual sensitivity scores when comparing the 3 major categories (1w-ANOVA, 3 levels; F_2,64_=58.590, p<0.001, η^2^=0.29). We have to note that this pattern of results is unlikely to be driven by the acoustic (dis-)similarity of some the 6 generic screams, as the machine-learning approach in the previous paragraph has shown a relative large acoustic distance (i.e. high above chance discrimination of the machine classifier; Fig. 1d) between all types of screams.

### Impaired perceptual discrimination of alarm compared to non-alarm screams

Experiment 2 showed that the 6 generic scream calls as well as the three major categories of scream types (neutral, non-alarm, alarm) differed in speed, accuracy, and sensitivity of their categorization when all types of screams and choice options were available in a multi-option decision task. Especially alarm screams showed the least neurocognitive processing efficiency (i.e. reaction time measures) and perceptual sensitivity (i.e. d’ measure) compared to non-alarm screams. This surprising pattern of higher processing efficiency of non-alarm than alarm screams might be specific to the more complex 7-alternative forced-choice task of experiment 2. This pattern might be different in simpler discrimination tasks while classifying only a low number of possible scream types. Specifically, in daily life, individuals often only need to choose only between 2 scream type options depending on certain contexts. In threatening situations, for example, individuals need to discriminate fear from anger, but not from other types of screams.

In experiment 3, another sample of participants (n=35) therefore discriminated screams in any combination of only 2 types of screams presented in the same block (e.g. joyful and fearful screams), with 21 possible combinations each using a simple two-option decision task (e.g. classify screams as joyful or fearful) (Fig. 2e-h). We found that only some combinations of screams led to differences in reaction times and accuracy between the two types of screams presented. Participants responded to positive pleasure screams (t-tests, all FDR corrected; t_34_=3.022, p=0.033) and joy (t_34_=4.858, p<0.001) faster when these screams were presented together and compared with fear screams in one block, as well as to positive pleasure screams when presented together and compared with sad screams (t_34_=3.730, p=0.007) in one block (Fig. 2d-e). In addition, more discrimination errors occurred for all negative screams (i.e. screams with a negative affective valence) when presented together with neutral screams (sad: t_34_=3.358, p=0.005; pain: t_34_=4.980, p<0.001; fear: t_34_=3.435, p=0.005; anger: t_34_=3.303, p=0.005) or with joyful screams (sad: t_34_=7.615, p<0.001; pain: t_34_=6.441, p<0.001; fear: t_34_=7.213, p<0.001; anger: t_34_=7.973, p<0.001); for angry screams when presented together with pleasure screams (t_34_=2.652, p=0.025); and for neutral screams when presented together with joyful screams (t_34_=3.787, p=0.002) (Fig. 2f-g). This pattern of results suggests that, compared to alarm screams, non-alarm and positive screams again are processed more efficiently in terms of faster reaction times and discrimination accuracy in simpler two-choice decision tasks. There might be the possibility that alarm screams do not need to be discriminated very quickly, because the only need to activity the perceiving organisms to indiscriminately respond to any potential threat. While this could explain the increased classification times for alarm screams (i.e. respond first, and then discriminate), it does not explain the higher error rates, which still point to a classification and discrimination disadvantage for alarm screams.

Given the general distinction of these neutral, non-alarm, and alarm screams, we also asked in experiment 3 whether scream discriminations within and across these broader categories would influence participants’ performance. We found that discrimination of the 6 generic screams from neutral screams led to the overall fastest responses (1w-ANOVA; 4 levels, referring to the 4 possible combinations: neutral and screams, within non-alarm, within alarm as well as alarm with non-alarm; F_1,3_=56.449, p<0.001, η^2^=0.13) (Fig. 2d) and the highest accuracy rate (1w-ANOVA; 4 levels; F_1,3_=149.590, p<0.001, η^2^=0.44) (Fig. 2f) compared with all other combinations (i.e. with non-alarm combinations, within alarm combinations, alarm and non-alarm combinations). The reaction time increased and the accuracy rate decreased starting from discrimination within non-alarm screams to discrimination of non-alarm from alarm screams and finally to discrimination within alarm screams. Discrimination among alarm screams required significantly increased reaction times (post-hoc planned comparisons; all p’s<0.001) and had significantly decreased accuracy compared with all other combinations (all p’s<0.001). Again, we have to note that all types of screams were acoustically very distinct from each other given the high above chance level classification according to the outcomes from the machine learning approach (Fig. 1d), such that discrimination impairments between screams were unlikely driven by potential acoustic similarities.

### Neural efficiency and significance for non-alarm scream processing

With the observation from experiment 2 and 3 that alarm screams are processed with significantly lower neurocognitive efficiency as one would assume given their threat signaling nature, we performed a last experiment (experiment 4) in which we recorded the brain activity of another sample of human listeners (n=30) by using functional magnetic resonance imaging. We assumed that the human brain might process alarm screams with significant efficiency compared with non-alarm screams as quantified by the level of neural signals in [1] and the connectivity between brain areas that are central to affective sound processing [22,27]. Voice signals and affect bursts are usually processed in a distributed brain network, consisting of the auditory cortex, the amygdala, and the inferior frontal cortex (IFC) [1,22,28], which provide an acoustic and socio-affective analysis of these signals [29,30], respectively. To identify the neural dynamics of scream call processing in this network, we asked humans to listen to the same selected 84 screams as in experiments 2 and 3. During listening to these scream calls, participants performed a largely orthogonal gender decision task on the screams, with orthogonal meaning that the task is largely intendent from the emotional quality of screams. The task was introduced to maintain the attention of the participants to the experiment, and is often referred to as implicit but still strong processing of the affective quality of the stimuli that leads to consistent brain activations [31,32]. Reactions times did not differ between all 7 scream types (F_6,174_=2.371, p=0.094) nor between the 3 major categories of neutral, non-alarm, and alarm screams (F_2,58_=0.756, p=0.428). The error rate was different between all 7 scream types (F_6,174_=7.280, p<0.001, η^2^=0.10), but not between the 3 major categories (F_2,58_=2.177, p=0.132) (Fig. S2).

While alarm screams (pain, anger, fear) mainly elicited lower neural activity in many inferior frontal and high-level auditory cortex regions, non-alarm screams (pleasure, sad, joy) compared with neutral vocalizations showed higher and extended auditory cortical activations, especially in the right hemisphere in the low- and high-level auditory cortex (Fig. 3b; Fig. S3, Tab. S1-2). This pattern of neural activity is enhanced when the perceived valence of the screams is taken into account (Fig. S4). The activation in the auditory cortex largely extended across the voice-sensitive cortex, as determined by a separate voice localizer scan (Fig. 3a; methods). When we compared neural activity for alarming screams with that of non-alarming screams (Fig. 3c), the non-alarming screams revealed extensive higher activity that was largely extended over the auditory and inferior frontal cortex. This indicates that a lower, not a higher, alarm level of screams is able to elicit more activity across many auditory and frontal brain regions; this was also shown in a separate analysis using the alarm rating across all screams as a parametric repressor, independent of the scream class (Fig. 4). Furthermore, neural activity for positive (pleasure, joy) as opposed to that for negative screams (sad, pain, fear, anger) elicited similar largely extended activity in the auditory and inferior frontal cortex, as well as some activity in the bilateral amygdala (Fig. 3d). Additional activity was found in the right posterior superior temporal sulcus (pSTS) for positive versus neutral vocalizations (Fig. 3c).

**Figure 3.**
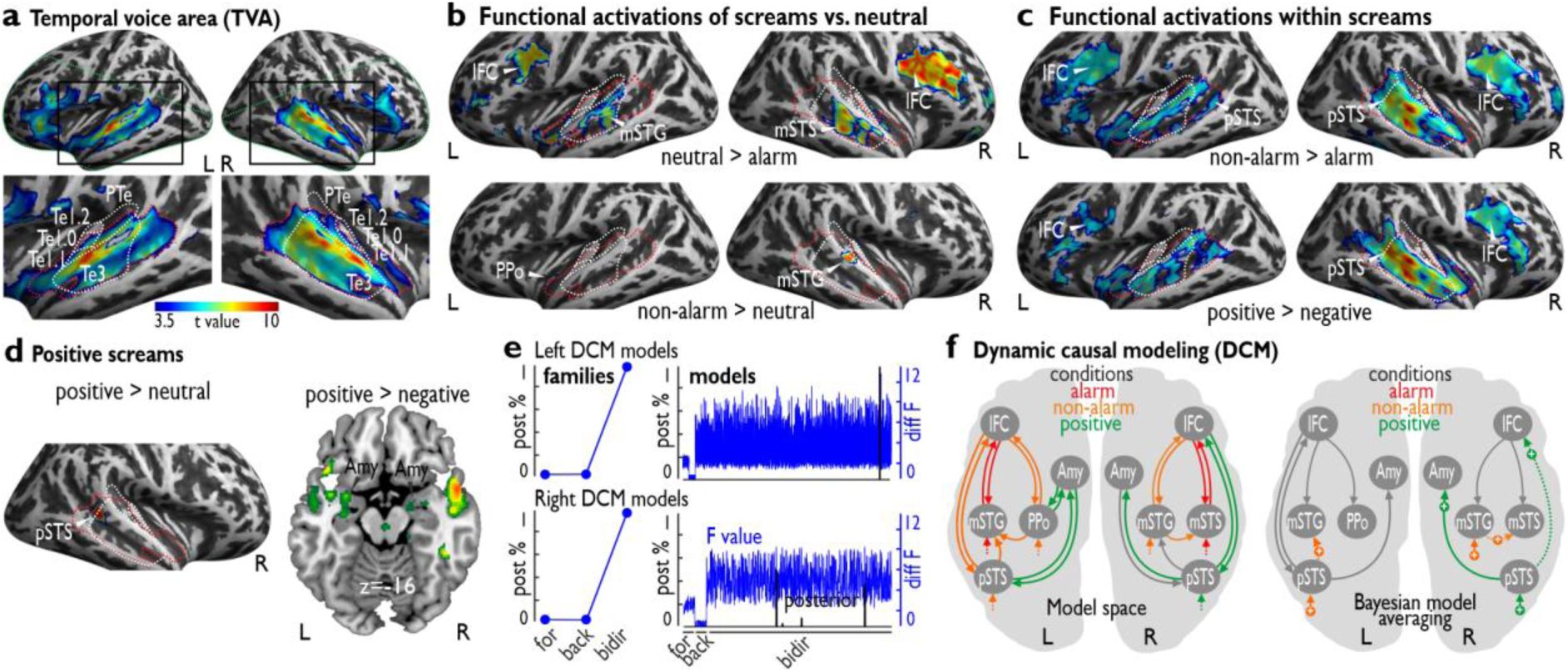
Neural activity and effective functional network for scream call processing. (a) Functional definition of the temporal voice area (TVA). The red dashed line indicates the cortical extension of the TVA; the green dashed line indicates the coverage of the partial volume acquisition; the white dashed line defines the anatomical subregions of the auditory cortex (primary areas Te1.0, Te1.1, Te1.2, and secondary area Te3) and the planum temporale (PTe). L = left; R = right. (b) Alarm screams compared with neutral screams elicited reduced activity in the bilateral IFC and STC (left mSTG, right mSTS), whereas non-alarm screams elicited increased activity in the bilateral STC (left PPo, right mSTG). IFC = inferior frontal cortex; STC = superior temporal cortex; mSTG = middle superior temporal gyrus; mSTS = middle superior temporal sulcus; PPo = planum polare. (c) Compared with alarm screams, non-alarm screams elicited significantly higher and extended activity in the bilateral IFC and STC. pSTS = posterior superior temporal sulcus. (d) Positive screams (pleasure, joy) showed higher activity in the right pSTS compared with that for neutral screams and in the bilateral amygdala compared with that for negative screams (sad, pain, fear, anger). Amy = amygdala. (e) Dynamic causal modeling (DCM) revealed the bidirectional model family as the winner family based on the posterior probability (left panel) in both the left (top left) and right hemispheres (bottom left). The right panel shows the posterior probabilities (black) and the log-evidence (blue; “diff F” = F-value minus the minimum F-value across models) for each model from the forward (for), backward (back), and bidirectional model families (bidir). (f) Bayesian model averaging (BMA) in the bidirectional model family indicated specific significant modulation of the left and right hemispheres only in the non-alarm (orange) and positive scream conditions (green). All activations are thresholded at p=0.005 voxel level and a minimum cluster size of k=42, resulting in a corrected threshold of p=0.05 at the cluster level.

**Figure 4.**
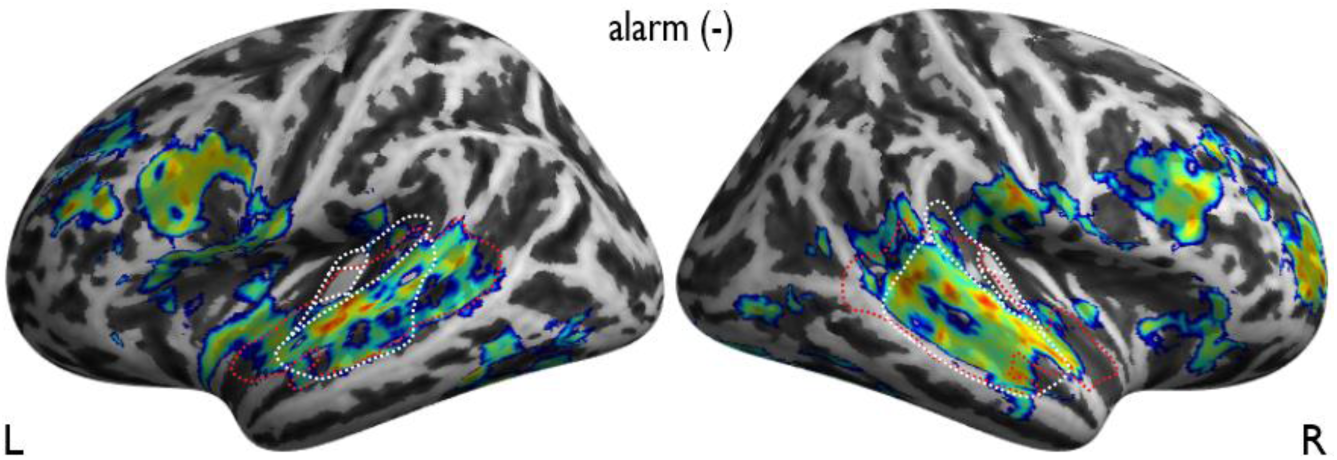
Negative parametric modulation of brain activity by the alarm rating of scream calls. While we did not find neural activity for a positive relationship with the alarm ratings of every scream, we found largely extended activity in the auditory cortex and the frontal cortex for negative associations with the alarm level of each scream. Threshold p=0.005 voxel level, cluster size of k=42 (corrected p=0.05 at cluster level).

To further examine the neural sensitivity for non-alarm screams, we performed an effective functional connectivity analysis on the observed patterns of brain activations using a dynamic causal modelling (DCM) approach. We sought to determine the neural information flow underlying the processing of alarm, non-alarm, and positive screams between the auditory cortex (sensory input and analysis), the IFC (cognitive evaluation), and the amygdala (affective analysis) in the left and right brain hemispheres (Fig. S5, Tab. S3). Determining this neural information flow can provide a detailed picture about the integrated functioning and directed interaction of diverse brain systems for decoding socio-biologically relevant information on sensory, cognitive, and affective levels. Using DCM (Fig. 3e-f), we first found a bilateral basic (intrinsic) neural network with top-down and bottom-up connections from and to the frontal from auditory cortical subregions, respectively, as well as intrinsic connectivity between the posterior auditory cortex (pSTS) and the amygdala. This intrinsic network was independent of any experimental condition. Second, in terms of experimentally driven modulations in this network, we found significant driving input effects of positive screams in the right hemisphere, especially of non-alarm screams in both hemispheres, to the auditory cortical subregions. Third, non-alarm screams also modulated specific connections between auditory cortical subregions in both hemispheres, while positive screams modulated the connectivity from the posterior auditory cortex to the amygdala and the frontal cortex in the right hemisphere. No input, intrinsic, or modulatory connectivity effects were found for alarm screams in both hemispheres.

## DISCUSSION

Our data provide evidence for a broad diversity of human scream calls with different affective meaning and communication profile. The data from experiment 1 aimed at a broad psychoacoustic description of human scream calls, and these data go beyond the frequent assumption that scream calls are of a generic alarming nature [1,5,8,19]. Instead of scream calls being of a uniform acoustic and communication nature, related to threat and alarm signaling based on fear, we found several distinctive scream categories of alarming, non-alarming, and even of positive nature in human primates. This distinctiveness was shown using a machine-learning approach including an extensive set of relevant acoustic voice features [25]. Based on this approach, human scream calls appeared to acoustically differ significantly from other nonverbal affect vocalizations in a cross-vocalization type of comparison, but they are also distinctive across different types of screams when compared within the set of the 6 generic screams. The acoustical patterns of screams thus seem to be distinct and largely specialized in the broad acoustic space of any kind of emotional vocalizations [6,21,22,26].

Besides this approach to compare scream calls based on a large acoustic feature set (i.e. 88 acoustic features), we also quantified another complex acoustic feature related to the spectral and temporal modulation rate of the acoustic signal and is referred to as modulation power spectrum (MPS) [23,24]. We found that the spectral modulation rate showed high power across a broad frequency range, but the temporal modulation rates were not distributed across a broad frequency range (30-150Hz) as previously reported [1]. Instead, we found that the temporal modulation rate was focused around two separate frequency bands centered on ∼60Hz and ∼160Hz, contributing to the harsh and rough acoustic quality of scream calls [33]. Furthermore, we also found that this roughness feature was not only characteristic of fearful scream calls as part of their alarming quality [1,19], but also of other alarm screams (pain, anger) and, most importantly, of some non-alarm screams (pleasure, sad). Thus, while acoustic roughness seems to be an important feature of many scream calls, it is also critically related to the perceived alarm level of scream calls especially centered on the low temporal frequency band of ∼60Hz given a higher beta weight in the linear regression analysis for this frequency band. The temporal modulation rate of ∼60Hz and its range from 50-70Hz corresponds to the transition zone between acoustic flutter and a more continuous but rough pitch perception of sounds [34,35] with a potential sensitivity of the (primary) auditory cortex to such features [34,36,37].

In experiment 1 we demonstrated the psychoacoustic distinctiveness of various types of human scream calls being both of alarming and non-alarming nature. In experiments 2 and 3 we then turned to the question of the decisional processes that are involved while categorizing (i.e. into the 6 plus 1 scream categories) and discriminating different types of screams (i.e. discriminating two types of screams), and especially of alarm and non-alarm screams. Contrary to an assumed higher neurocognitive efficiency in processing alarm screams based on their proposed sociobiological relevance [1], alarm scream calls were overall responded to the slowest, had generally lower categorization accuracy, and were associated with a lower perceptual sensitivity, and were finally discriminated the worst in a two-class decision task. There were some exceptions to this general observation that alarm scream were classified slower and with higher error rates. For example, sad screams were classified slower when specifically compared to anger screams; furthermore, the performance for joy screams had large effects on the mean performance during non-alarm scream classifications. But when quantifying the d prime measure as an indicator of perceptual sensitivity as well as the false alarm rate, the general observation of a lower processing efficiency of alarm compared to non-alarm scream was largely confirmed. These findings are very surprising given that speed and accuracy in processing as well as a perceptual sensitivity of the primate neurocognitive system is frequently assumed to be crucial for signals that indicate threat and danger [1,31,38–40].

This impairment in efficiently classifying alarm screams with the exceptions mentioned above was not only found in the more complex multi-option decision task of experiment 2, but also in the simpler two-option decision task of experiment 3. We have to note that we used the exact same stimulus material in experiment 2 and 3, but critically changed the decisional tasks that participants had to perform. In experiment 3 we asked participants to only decide between two possible scream types, and found the highest reaction times and error rates while discriminating between alarm scream. The latter results seem contrary to the usual assumption that discrimination of alarm screams, among themselves and from non-alarm screams, demands fast behavioral response and differential coping strategies since we found the opposite pattern, such that alarm screams take the longest to respond to and were discriminated with the lowest accuracy. This additionally suggests that alarm screams, surprisingly, need more processing effort during simple scream discrimination with only two choice options, and that rather non-alarm and positive screams are more efficiently processed during this perceptual discrimination of two types of screams. Non-alarm screams and positive screams might be more relevant in social environments and interactions of human beings, and also might show a higher occurrence rate in daily lives [41]. The classification and discrimination of non-alarm and positive screams might thus have priority for humans and might explain their better processing efficiency. The only priority that alarming scream categories received was their frequent use during misclassifications of other screams in a multi-option classification task in experiment 2. This might resemble a natural threat perception bias that seems to be a cost-benefit efficient solution when balancing response options against potential sources of (non-)threat, especially under conditions of uncertainty [42]. One further specific note concerns the various combinations of two specific scream types in experiment 3. Some of these combinations might only rarely occur in daily life, but for completeness we include any possible combination here. For example, specific combinations of non-alarm and alarm screams might occur less frequently in daily life (e.g. joy and anger screams), but there nonetheless certain contexts that include co-occurrences of non-alarm and alarm screams (e.g. a happy social gathering, where an interactions between two people suddenly turns violent because of an incident).

In accordance with a decisional impairment when classifying and discriminating alarm compared to non-alarm screams, the neural patterns of functional activations in experiment 4 pointed to a sensitivity of the human neural system for non-alarm rather than for alarm screams in the neural system. The neural processing of alarm screams largely resulted in decreased neural activations in comparison to other types of screams. Non-alarm and positive screams elicited increased neural activations in brain regions such as the inferior frontal cortex, the auditory cortex, and the amygdala. These regions are a core neural network that support the social evaluation [43], acoustic analysis [27,29], and affective assessment of sounds [44,45]. Especially the auditory cortex and the amygdala were previously assumed to be reserved only for negative and alarming voice signals [1,27,46]. Using a parametric analysis approach we also found that low, rather than high, levels of the alarming quality of screams were mainly driving the neural activity that we similarly found for comparing non-alarm and positive screams against other types of screams. Thus, the neural system seems not to be directly sensitive to detect alarm signals, at least when encoded in scream calls, but seems to be more sensitive to non-alarm and positive emotions. These neural effects resemble the behavioral decision-making profiles that we found in experiment 2 and 3.

Furthermore, modelling the directional neural connectivity between the frontal, auditory, and limbic brain regions, we also found that non-alarm screams, especially positive screams, show significant effects in neural network dynamics over alarm screams, such that the human neural system is more prepared to decode non-alarming and positive information signaled in human scream calls in a mostly bottom-up manner. These findings in humans are surprising and largely diverge from studies in non-human primates; for latter, the literature so far only reported scream calls being expressed in negative contexts. While screams can also be expressed in negative contexts in humans, humans seem to be the only species to express screams in non-alarming and especially in positive contexts. Concerning these non-alarming contexts and our neural connectivity analysis, we found that non-alarm screams provided significant input modulation to some auditory cortical subregions in both hemispheres and a modulation of a connection of auditory subregions in the right auditory cortex, positive screams showed significant input modulation in the right hemisphere as well as a modulation of the forward connection from the right posterior auditory cortex to both the right amygdala and the right inferior frontal cortex. Thus, there seem to be specific neural pathways connecting subregions in the right auditory cortex for non-alarm scream processing as well as connecting the auditory cortex with the frontal cortex and the limbic system to specifically decode positive meanings from screams. Previous work has shown a connectivity from the auditory cortex to the amygdala [27] for acoustic information transfer to perform affective assessment, but only for unpleasant sounds. The auditory cortex also shows forward connection to the frontal cortex [47,48], but this forward connection was so far not shown to be modulated by the affective meaning of sounds.

Our data critically extend these previous findings, especially showing that these pathways in the right hemisphere are predominantly sensitive to positive voice calls. Most surprisingly again, alarm screams did not show any effects in this directional connectivity analysis either as driving input to specific brain regions, as intrinsic connectivity, or as modulation of the connections between regions. We have to note that all neural activations and specifically all connectivity results were obtained while participants performed a non-emotional gender decision task, and thus participants were not explicitly focusing on the affective quality of the screams. However, the implicit processing of the affective quality of voice signals has been shown by many previous studies [31,32], and given that the gender task was largely unrelated to the affective and alarming quality of the screams types, we think that the neural activations and connected patterns largely reflect the neural dynamics for generic scream processing and scream discrimination.

Taken together, this overall pattern of psychoacoustic, decision-making, and neural results for alarm screams in humans seems rather unusual for sociobiological voice calls when reviewed in the broader animal communication field. Scream calls were previously assumed to be most crucial in signaling and accurate communication of alarm across a broad range of animal species [5,8,10], sometimes also including references to humans [1,19]. Scream calls in non-human primates and other animals have been so far been reported to be expressed and perceived exclusively in negative contexts. A different picture of scream calls seems to emerge when investigated in humans, such that human listeners overall respond more quickly, more accurately, and with higher neural sensitivity to non-alarm and positive scream calls, which seem to have a higher relevance in human sociobiological interactions [49–51]. There seem some exceptions from this overall pattern of scream recognition in humans, but across many psychoacoustic, behavioral, perceptual, and neural effects quantified here, alarm screams often show less neurocognitive processing efficiency than non-alarm screams. Alarm cream categories only have some primacy during misclassification of other scream types, which might be a safety choice under conditions of decisional uncertainty. And this safety choice might be shared with other non-human species that use screams in their vocal repertoire.

## MATERIALS AND METHODS

### Participants

All four experiments included an independent sample of human participants with normal hearing abilities and normal or corrected-to-normal vision. No participant presented a neurological or psychiatric history. All participants gave informed and written consent for their participation in accordance with the ethical and data security guidelines of the University of Zurich. The experiments were approved by the cantonal ethics committee of the Swiss canton Zurich.

### Experiment 1 – Acoustic dimension of scream calls

#### Participants

In experiment 1, 12 healthy volunteers (6 males; mean age 29.08 years, SD=5.66, age range 22-42) took part in the acoustic recording of 6 different types of screams, as well as a recording of neutral vocalizations as an intensive utterance of the vowel /a/ as a seventh category.

#### Experimental design

We invited participants to vocalize 6 different types of screams. These types of screams were chosen to comprehensively cover all possible screams that humans vocalize in certain emotional states. A previous report on the acoustic and neural processing of screams identified screams as being largely only of a negative fearful nature [1] to signal alarm to other conspecifics [19] on the basis of a certain and unique acoustic feature of “roughness” (i.e. high-frequency spectro-temporal acoustic modulations). Although fearful screams are a prominent example of human screaming, humans produce vocal screams of a rough acoustic nature not only in the emotional state of fear, but also in a variety of emotional states referred to as “pleasure,” “sadness,” “joy,” “pain,” “fear,” and “anger” states. These different types of screams can be classified as either positive (pleasure, joy) or negative screams (sadness, pain, fear, anger), which we refer to as the factor “valence.” Furthermore, screams can be classified as either alarming vocal signals (pain, fear, anger) or as non-alarming vocal signals (pleasure, sadness, joy), which we refer to as the factor “alarming quality.” Thus, screams are not only limited to negative and alarming screams of fear, but they can also be positive and of a non-alarming nature.

We thus instructed the 12 participants to produce vocalizations of screams for each of the 6 types of generic screams, as well as to produce neutral screams on the basis of an intense vocalization of the vowel /a/. For each type of scream, we provided short instructions to each participant to imagine several corresponding context(s) in which these screams are commonly produced (e.g. fearful scream: “You are being attacked by an armed stranger in a dark alley”; anger screams: “You try to intimidate an opponent”; joyful screams: “Your favorite team wins the World Cup”; pleasure screams: “You are screaming from sexual delight”). For each participant, we recorded 8 instances of each type of scream in an anechoic chamber with a Neumann TLM-102 microphone at a distance of ∼1m from the speaker. From these 8 instances, we chose the 5 best recordings from perceptual judgment of recording quality, vocalization length (800-900ms), and continuous vocalization for the duration of the scream. The final selected screams were cropped to a fixed duration of 800ms, standardized to an identical RMS across screams corresponding to 70dB SPL, and faded-in and faded-out by a 15ms intensity ramp at the beginning and end of each scream. The final sample included 420 screams. Although this standardization of the scream to 800ms and 70 dB SPL might render some of the screams partly unnatural (i.e. alarming screams might have a higher natural loudness/intensity than non-alarm screams), this procedure ensured that all following analyses and perceptual experiments are not confounded by basic acoustic features of these screams due to vocalization characteristics and recoding conditions for single speakers. To avoid experimental confounds, we rated the exact standardization of the stimuli of higher importance for a straightforward interpretation of the stimuli compared to introducing a little bit less natural sounding screams. Furthermore, although these screams were largely acted rather than spontaneously expressed, acted screams seemed to be perceptually similar to natural screams [1,2], and thus provide a valid basis for affective communication research in human and nonhuman primates.

#### Perceptual assessment of screams

After the acoustic recording of the screams, first, we asked 23 participants (12 males; mean age 24.04 years, SD=5.35, age range 20-46) to perceptually assess the alarming quality of each of 420 screams on a continuous scale ranging from “0” (not alarming at all) to “1” (highly alarming). Second, we asked another sample of 26 participants (11 males; mean age 24.40 years, SD=3.56, age range 19-32) to perceptually assess these screams according to the central emotional dimensions of arousal and emotional category. Arousal was quantified on a 7-point Likert scale ranging from 1 (“not arousing at all”) to 7 (“highly arousing”); emotional category was quantified by a 7-alternative forced-choice (7AFC) classification (“neutral,” “pleasure,” “sad,” “joy,” “pain,” “fear,” “anger”). From this perceptual assessment and recognition accuracy, we selected 84 screams from 3 male and 3 female speakers (2 instances per scream type for each speaker) for the subsequent experiments (see below).

#### Data analysis

The data from the perceptual assessment were separately analyzed for the arousal ratings and the classification ratings. Arousal ratings were subjected to a repeated-measures 1-way ANOVA that included the within-subject factor scream type (levels: neutral, pleasure, sad, joy, pain, fear, anger). Using a similar ANOVA, we also analyzed the classification accuracy of the screams. In addition, we quantified the confusion matrix (i.e. individual percentage of categories chosen during misclassification of each type of scream) and the false alarm rate (i.e. general percentage of categories chosen during misclassifications). The significance threshold was set to p=0.05 (here and for all analyses that follow).

The alarm ratings were analyzed in a similar way as for arousal and valence ratings. For an additional analysis of the alarm ratings, we used a full permutation and statistical testing through any possible categorical combination of the 6 generic screams, with the described categorization leading to the most significant distinction in the alarm level between screams (see methods). During this permutation approach, one category was always represented by neutral screams, while the generic 6-scream types were randomly split into 2 separate categories of screams with 2-4 scream types per category. This led to 25 possible combinations of screams. For each combination, we calculated the mean alarm rating for each participant and subjected these data to a 1-way repeated-measures ANOVA with 3 levels.

#### Acoustic analysis

The acoustic analysis of screams and other vocalizations was based on 88 previously described acoustic features [25] extracted with the toolbox openSMILE [52]. The acoustic features set comprised frequency (pitch, jitter, etc.), energy/amplitude (intensity, shimmer, etc.), and spectrum-related features (alpha ratio, Hammarberg index, etc.), as well as 6 features related to the temporal domain (rate of loudness peaks, voiced segments, etc.). Arithmetic mean and coefficient of variation (standard deviation normalized by the arithmetic mean) were returned for all parameters. For the loudness parameter, the 20th, 50th, and 80th percentiles; the range of 20th to 80th percentiles; and the mean and standard deviation of the slope of rising/falling signal parts were also computed. Each acoustic parameter was normalized (i.e. centered on the mean and divided by its standard deviation) across all 420 sounds.

Besides these basic acoustic features, we also analyzed the modulation power spectrum (MPS) [24] by using the MPS toolbox for MATLAB (theunissen.berkeley.edu/Software.html). To obtain an MPS for each sound, we first converted the amplitude waveform to a log amplitude of its spectrogram obtained by using Gaussian windows and a log-frequency axis. The MPS results from the amplitude squared as a function of the Fourier pairs of the time (i.e. temporal modulation in Hz) and frequency axis (i.e. spectral modulation in cycles/octave) of the spectrogram. The low-pass filter boundaries of the modulation spectrum were set to 200Hz for the temporal modulation rate and to 12 cycles/octave for the spectral modulation rate. A statically difference of the MPS from generic scream compared to neutral screams was tested using permutation approach (n=2000) by shuffling scream type labels, resulting in a p-value map for the entire spectral and temporal range of the MPS (Fig. 1b).

#### Machine learning approach

Using the 88 acoustic features that were extracted from each scream call using the openSMILE toolbox as described above, we then used a machine learning approach to assess the acoustic distinctiveness of the various types of screams. For each generic scream category and the neutral vocalizations, we trained a support vector machine (SVM) [53] to discriminate these seven emotion categories of screams on the basis of all acoustic parameters as features (see [54] for a discussion of classification methods). This approach is called a “cross-validation” approach. The SVM is a binary classifier, and so multiclass classification can be set up in either a 1-vs-1 or a 1-vs-all scheme [55]. We used a 1-vs-1 scheme as implemented by MATLAB’s fitecoc function (MATLAB, version 2018a), which estimates a classification model for each available pair of vocalization categories separately. The SVM parameters were set to the MATLAB default values and the kernel function was a third-order polynomial.

To evaluate the classification accuracy for the scream discrimination in the cross-validation approach, we tested the SVM on unseen testing data that were not used to estimate the model parameters. Within each scream category, a 5-fold cross-validation scheme split the sound set (420 sounds, with 60 sounds per emotion) repeatedly into a training and a test data set, such that each sound served once as a test data point. This approach estimates how well the SVM is capable of separating the seven scream categories (6 generic screams, 1 neutral vocalizations) based on their acoustic features.

Next, a new SVM model learned to separate all scream types and predicted the emotion label on another set of more standard nonverbal expressions of emotion (e.g. joyful laughter and anger growls) of the same valence as the scream types and vice versa. The nonverbal vocal expression consisted of 448 emotional voices from 8 male and 8 female speakers (mean age 30.38 years, SD=7.07, age range 21-47; 10 of these 16 speakers also vocalized the scream calls used here) that included the vocal expressions neutral (vowel /a/ in a normal voice), pleasure, sad, joy, pain, fear, and anger. These nonverbal vocal expressions were recorded with the same kind of settings that we used to record the scream calls. From every speaker, we had 4 instances of a vocalization for each emotion. This approach is referred to as “cross-classification” approach, and tests whether a model learned a representation of the emotion categories that generalizes across vocalization categories (screams, nonverbal expressions).

### Experiment 2 – Perceptual categorization of scream calls

#### Participants

In experiment 2, Thirty-three healthy volunteers (12 males; mean age 23.91 years, SD=3.28, age range 19-33) took part in the experiment on the perceptual categorization of 7 different types of screams, including the seventh neutral vocalization.

#### Experimental design

From the acoustic scream recordings of experiment 1, we selected 84 screams (3 male, 3 female speakers) with 2 instances of screams per category. Stimuli were selected from the results of the perceptual assessment of screams in experiment 1, such that no significant differences in the recognition rate across scream types (F_1,6_=1.895, p=0.085) were found for this selection. Mean arousal level differed across all 7 scream types (F_1,6_=51.065, p<0.001), across the 6 generic screams (F_1,5_=11.647, p<0.001), and between neutral, alarm, and non-alarm screams (F_1,2_=77.645, p<0.001). The experiment consisted of 3 sessions. One session included 168 trials and, during each session, the 84 screams were presented twice. Screams were presented at 70dB SPL using how quality headphones with a flat frequency profile (Sennheiser® HAD 280), and participants were asked to perform a 7AFC task according to the emotional valence of the screams (“neutral,” “pleasure,” “sad,” “joy,” “pain,” “fear,” “anger”). Decision options were presented on the screen in a circular arrangement, each at ∼5° visual angle from the center of the screen. Participants had to respond by moving the mouse cursor from the middle of the circle (i.e. mouse cursor position was reset to the middle of the circle after each trial) to one of the response options and pressing the left mouse button with the right index finger. Participants were asked to respond as quickly as possible, but still trying to make an accurate decision. The arrangement of response options was randomly chosen for each participant but was kept constant across blocks for each participant. Screams were presented in random order at 70dB SPL, with a maximum response time window of 3000ms; trials with no response in this time window were classified as misses trials (6.53% of all trials). After a response option was chosen, the response screen disappeared and the next scream was presented after an inter-trial interval (ITI) of 1750 +/- 250ms.

#### Data analysis

Reaction time (RT) and classification accuracy data were separately subjected to a repeated-measures 1-way ANOVA that included the within-subject factor scream type (levels: neutral, pleasure, sad, joy, pain, fear, anger). RTs were defined as the response latency from stimulus onset to the button press, and RTs were only calculated for correct trials. Accuracy was defined as pressing the one button that corresponded to the pre-assigned category of a scream (e.g. pressing the button for “fear” when a fear scream was presented). We performed an additional 1-way ANOVA that included 3 categories: neutral, alarm, and non-alarm screams. We again quantified the confusion matrix (i.e. individual percentage of categories chosen during misclassification of each type of scream) and the false alarm rate (i.e. general percentage of categories chosen during misclassifications). From the correct classification for a scream category and the level of false alarm rates that used a certain scream category, we calculated the d′ measure as an indicator of the sensitivity and detectability of a certain scream type. The d′ measures were subjected to the same ANOVA analyses as described above.

### Experiment 3 – Perceptual discrimination of scream calls

#### Participants

In experiment 3, Thirty-five healthy volunteers (11 males; mean age 24.34 years, SD=3.25, age range 20-30) took part in the experiment on the perceptual discrimination of screams.

#### Experimental design

The same 84 screams as in experiment 2 were used. Screams were presented in 21 blocks, each block containing 2 different scream types. Over the 21 blocks, we presented all possible combinations of the 7 scream types. One block consisted of 72 trials. For example, in one block, we presented only angry (36 trials) and fearful screams (36 trials). Screams were presented as single trials in a random order, and after each scream participants performed a 2-alternative forced-choice (2AFC) task by pressing one button (left index finger) for one scream category (e.g. “anger”) and another button (right index finger) for another scream category (e.g. “fear”). Participants were again asked to respond as quickly as possible, but still trying to make an accurate decision. Trials with no response in this time window were classified as missed trials (1.55% of all trials). In each block, screams were presented in random order at 70dB SPL, with an ITI of 2250ms +/-250ms. Button and response option assignments were counterbalanced across blocks and participants.

#### Data analysis

For each of the 21 combinations of screams, we scored the mean RTs (i.e. response latency from stimulus onset to button press, only for correct trials) and accuracy data (i.e. correct trials were defined as button press that corresponded to the pre-assigned scream type) for each of the 2 scream categories, as well as the mean difference in RTs and accuracy data between the 2 categories for each participant. The mean difference values for each scream combination were normalized by using the mean RT (1/mean RT) or the mean accuracy rate (mean %) for the 2 scream types for each participant. For each of the 21 scream combinations, we tested for significant differences in RTs and accuracy data by using a t-test; the significance threshold was adjusted for multiple testing by using the false discovery rate (FDR) correction. We performed an additional 1-way ANOVA by comparing mean RTs and accuracy rates for 4 categories of potential scream combinations: neutral screams combined with generic screams (e.g. neutral combined with fear screams), non-alarm screams combined with non-alarm screams (e.g. pleasure combined with sad screams), non-alarm screams combined with alarm screams (e.g. pleasure combined with anger screams), and alarm screams combined with alarm screams (e.g. fear screams combined with anger screams).

### Experiment 4 – Neural processing of scream calls

#### Participants

In experiment 4, *w*e invited 30 healthy volunteers to take part in this experiment (15 males; mean age 26.10 years, SD=4.95, age range 18-43). Participants were informed about the aim to investigate the neural dynamics of processing different types of screams, and they were informed to perform a task that is orthogonal to the emotional dimension of screams.

#### Experimental design

During the functional magnetic resonance imaging (fMRI) experiment, we presented the same 84 screams as in experiment 2. The experiment included 2 blocks, and each block included 168 trials. Screams were presented in random order at 70dB SPL, with an ITI of 3875ms +/-875ms. For the presentation of the stimuli, we used active noise-cancellation MR-compatible headphones (OptoACTIVE II; www.optoacoustics.com) that can reduce MRI scanner noise by up to 20dB. Participants performed a gender decision task, which is a task that is orthogonal to the relevant dimension of processing the different types of screams. This gender task has been used in previous studies and was established to be validly orthogonal to the emotion dimension of stimuli [28,31]. After each scream presentation, participants indicated the corresponding gender of the speaker by pressing one button (right index finger) when the speaker was “male” or the other button (right middle finger) when the speaker was “female”. Trials with no response in this time window were classified as missed trials (1.87% of all trials). This basic task was implemented to keep the participants attentive throughout the experiment. Participants were again asked to respond as quickly as possible, but still trying to make an accurate decision. The task was orthogonal to the affective valence dimension of screams as the target dimension for the main analysis focusing on the affective valence of screams. Response button assignment was counterbalanced across participants but was constant across blocks for a single participant. Functional brain data were recorded on a on a 3T-Philips Ingenia by using a standard 32-channel head coil. Data pre-processing was done according to standard principles including a normalization to the Montreal Neurological Institute (MNI) stereotactic space.

For the statistical analysis of the functional brain data, each experimental condition was modeled as a separate regressor in a subject-wise GLM analysis, and the resulting images were taken to a second-level group analysis. All functional group results were thresholded at a combined voxel threshold of p<0.005 corrected for multiple comparisons at a cluster level of k=42. Modelling the effective connectivity between activated brain areas resulting from the various contrasts performed was done using dynamic causal modelling (DCM) as implemented in SPM12 [56], and including a step-wise procedure (methods). We first created 3 different families of models (forward, backward, and bi-directional models) for each hemisphere separately on the basis of the definition of the input to regions (C matrix), the intrinsic connection between regions (A matrix), and the modulation of connection by experimental conditions (B matrix) (Fig. S4). After estimating the winning model family in each hemisphere, we secondly used a Bayesian model averaging (BMA) approach, which creates a weighted average of all models within the winning family by taking the model evidence of each model into account. The resulting posterior estimates for each parameter in the A, B, and C matrix and the weighted average model were tested for significance by using a t-test against “0,” with resulting p-values adjusted for multiple comparisons using the FDR correction.

#### Voice localizer scan

The stimulus material consisted of 500ms sound clips[57] consisting of 70 human speech and non-speech vocalizations and of 70 non-human vocalizations and sounds (animal vocalizations, artificial sounds, natural sounds) presented at 70dB SPL. Each sound was preceded by a 500ms fixation cross and followed by a jittered blank 3550 to 5000ms gap before the onset of the next stimulus. Each sound clip was presented once during the experiment. Participants were asked to perform a 2AFC task (right hand: index and middle finger; button assignment counterbalanced across participants) on the stimuli to decide whether they heard a vocal or a non-vocal sound. The experiment was preceded by 10 training trials (random selection of 5 vocal and 5 non-vocal trials) to familiarize the participants with the task. The training trials were discarded from further analyses.

#### Image acquisition

The recording of structural and functional brain data was done on a 3T-Philips Ingenia by using a standard 32-channel head coil. High-resolution structural MRI was acquired by using T1-weighted scans (TR 7.91s, TE 3.71ms, voxel size 0.57mm^3^; in-plane resolution 256×251). Functional whole-brain images were recorded with a T2*-weighted echo-planar pulse sequence (TR 1.65s, TE 30ms, FA 88°; in-plane resolution 128×128 voxels, voxel size 1.71×1.71×3.5mm; gap 0.4 mm; 19 slices) with particle volume acquisition of slices rotated ∼30° nose-up to the anterior commissure (AC)-posterior commissure (PC) plane. The partial volume covered the superior temporal cortex, inferior frontal cortex, and medial limbic system (i.e. the amygdala). Data pre-processing was done according to standard pipeline including a normalization to the Montreal Neurological Institute (MNI) stereotactic space and isotropic smoothing (6mm) of the functional data (methods).

#### Data preprocessing

Preprocessing and statistical analyses of functional images were performed with Statistical Parametric Mapping software (SPM12, Welcome Department of Cognitive Neurology, London; www.fil.ion.ucl.ac.uk/spm). Functional data were first manually realigned to the AC-PC axis, followed by motion correction of the functional images. Each participant’s structural image was co-registered to the mean functional image and then segmented to allow estimation of normalization parameters. Using the resulting parameters, we spatially normalized the anatomical and functional images to the Montreal Neurological Institute (MNI) stereotactic space. The functional images for the main experiment were resampled into 1.7mm^3^ voxels. All functional images were spatially smoothed with a 6mm full-width half-maximum isotropic Gaussian kernel.

#### Single-subject and group analysis

For the first-level analysis of the main experiment and the voice localizer scan, we used a general linear model (GLM), and all trials were modeled with a stick function aligned to the onset of each stimulus, which was then convolved with a standard hemodynamic response function. For the main experiment, trials were modeled for each of the 7 scream types by separate regressors in one GLM, and for each trial, we entered the arousal level of each scream from the pre-evaluation results into the GLM as a covariate of no interest. The scream types differ in their arousal level, which might confound the neural brain data. For the voice localizer scan, we modeled trials with vocal and non-vocal sounds separately in another GLM. For each GLM of the main experiment and the voice localizer scan, we also included 6 motion correction parameters as regressors of no interest to account for signal changes not related to the conditions of interest.

Contrast images for the main experiment were then taken to a random-effects group-level analysis to investigate the neural processing of the different types of alarming and non-alarming screams by performing several directed contrasts between conditions (Fig. S2). Contrast images for the voice localizer scan were then taken to separate random-effects group-level analyses in order to determine voice-sensitive regions in both hemispheres of the auditory cortex. All group results were thresholded at a combined voxel threshold of p<0.005 corrected for multiple comparisons at a cluster level of k=42. This combined voxel and cluster threshold corresponds to p=0.05 corrected at the cluster level and was determined by the 3DClustSim algorithm implemented in the AFNI software (afni.nimh.nih.gov/afni; version AFNI_18.3.01; including the new (spatial) autocorrelation function (ACF) extension) according to the estimated smoothness of the data across all contrasts. The cluster extent threshold of k=42 was the maximum value for the minimum cluster size across contrasts of the main experiment and the functional voice localizer scans.

For the main experiment, after the GLM analysis with predefined scream categories based on the original scream recordings and scream selection (Fig. 3), we performed the same analysis by using the subjective post-experimental scream classification of each participant (Fig. S3). In this analysis, trials for the 7 scream categories were defined from the classification of each participant; e.g. an original anger scream that was classified as a “sad” scream by a participant was used as a sad scream trial in the analysis. Thus, this analysis was based on the perceptual impression of each participant rather by the preassigned scream classification. This analysis must, however, be interpreted with caution. First, it resulted in an unequal number of trials for each category and for each participant, since this depended on the classification performance of each participant. Second, the analysis is not sensitive to the classification errors of participants. Some of the trials might thus include error trials, where participants subjectively pressed the wrong button but recognized their error. Third, these scream classifications were performed after the experiment and not during the fMRI experiment. However, overall, this analysis might give a good estimate of the neural activity that underlies the perceptual impression of the participants.

We performed an additional parametric modulation analysis by modeling each trial on a single subject level by using the alarm ratings for each scream as a covariate, including one regressor for all trials and an additional regressor for the covariate in a GLM analysis. The arousal level was again included as a covariate of no interest. Single subject beta images were taken to a second-level group analysis as described earlier. Contrasts were computed for positive and negative associations with the alarm ratings as covariates of no interest (Fig. S4).

#### Dynamic causal modeling (DCM) of effective brain connectivity

We used the DCM12 toolbox implemented in SPM12 to model the information flow in the left and right brain hemispheres during the processing of different types of screams. We defined a set of regions of interest (ROIs) in the auditory cortex, amygdala, and IFC on the basis of group peak activations for the different contrasts performed. For each definition of a peak, the contrast was chosen that was the most informative about the potential sensitivity of the region for certain experimental conditions and scream types. The amygdala peaks (left [-29 2 -17], right [30 -3 -17]) were based on the contrast [positive > negative], and the IFC peaks (left [-44 10 29], right [42 9 27]) were based on the contrast [non-alarm > alarming]. In the left hemisphere, we furthermore defined peaks in the planum polare (PPo; [-42 - 7 -21], contrast [non-alarm > neutral]), middle superior temporal gyrus (mSTG; [-61 -25 5], contrast [neutral > alarm]), and posterior superior temporal sulcus (pSTS; [-63 --35 8], contrast [non-alarm > alarm]). In the right hemisphere, we defined the pSTS ([47 -37 5], contrast [positive > neutral]), mSTG ([63 -13 0], contrast [non-alarm > neutral]), and mid STS (mSTS, [54 -24 -2], contrast [neutral > alarm]). Signal time courses from these ROIs were extracted for each participant in a sphere with a radius of 3.4mm, including only supra-threshold voxels (p<0.05). The signal time course was quantified as the first eigenvariate representing the most typical time course across voxels included in each ROI. Acquisition delay for each ROI was accounted for by estimating the time of signal acquisition in each ROI across the time interval of the TR.

We created 3 different families of models for each hemisphere separately on the basis of the definition of the input to regions (C matrix), the intrinsic connection between regions (A matrix), and the modulation of connection by experimental conditions (B matrix). The first model family is referred to as “forward models” (n=128 models for left and n=64 models for right hemisphere) from the definition of the A matrix, including the bidirectional connections between regions in the auditory cortex, but only forward connections to the amygdala and the IFC. The family of “backward models” (n=128 for left and n=64 for right) is identical to the forward models, but only includes a backward connection from the amygdala and the IFC to the STC regions. The “bidirect models” (n=4096 for left and n=1028 for right) include bidirectional connections between the amygdala and the IFC on the one side, and the STC regions on the other side. For the left hemisphere, the input C matrix included non-alarm trials to the pSTS and the PPo, as well as alarm trials to the mSTG, while for the right hemisphere, the C matrix included positive trials to the pSTS, non-alarm trials to the mSTG, and alarm trials to the mSTS. This input C matrix was fixed for all models and only included subregions of the auditory cortex as potential regions for experimental input, but we estimated the strength and significance of these driving inputs for each model. Concerning the B matrix, we defined connectivity modulations from the predominant response of the seed and target region; for example, the connectivity to and from the amygdala was supposed to be only modulated for positive screams, since activation in the amygdala was mainly driven by those screams. For details about the B matrix specification, as well as the specification of the A and C matrix, see Fig. S4.

For each model family and each hemisphere, we estimated DCM for any possible combination of parameters in the B matrix, ranging from no modulation of connectivity to full modulation of any connection as defined in the A matrix. For the bidirect model family in the left hemisphere, this, for example, resulted in a total of n=4096 models. To define the winning model in each hemisphere, we used a 2-stage approach. First, we compared model families within each hemisphere and defined the winning family on the basis of Bayesian model selection by using a fixed-effect approach, assuming the same model architecture across the participants [58], which is a similar approach to that in a previous study on auditory processing [59]. Second, within the winning family, we used a Bayesian model averaging (BMA) approach, which creates a weighted average of all models within the winning family by taking the model evidence of each model into account. The resulting posterior estimates for each parameter in the A, B, and C matrix and the weighted average model were tested for significance by using a t-test against “0,” with resulting p-values adjusted for multiple comparisons using the FDR correction. For DCM in the left hemisphere, we had to exclude n=2 participants, and for right hemispheric DCM, we had to exclude n=2 participants, given extreme outlier values for several of the estimated parameters [59].

#### Post-experimental perceptual assessment of screams

After the fMRI experiment, we asked every participant to perform a 7AFC task according to the emotional valence of the screams (“neutral,” “pleasure,” “sad,” “joy,” “pain,” “fear,” “anger”), similar to that for experiment 2. This was done outside the scanner because this rather complex categorization task was difficult to implement inside the fMRI environment with 7 response options. Next to this valence classification, we asked participants to rate the arousal level of each scream beforehand on a 7-point Likert scale ranging from 1 (“not arousing at all”) to 7 (“highly arousing”).

Classification accuracy data were separately subjected to a repeated-measures 1-way ANOVA that included the within-subject factor scream type (levels: neutral, pleasure, sad, joy, pain, fear, anger). We again quantified the confusion matrix (i.e. individual percentage of categories chosen during misclassification of each type of scream) and the false alarm rate (i.e. general percentage of categories chosen during misclassifications).

## ACKNOWLEDGMENTS

This study was supported by the Swiss National Science Foundation (SNSF PP00P1_157409/1 and PP00P1_183711/1 to SF). All authors declare no conflict of interests. We thank Plamina Dimanova for help during data acquisition, and we thank Claudia Roswandowitz, Simon Townsend, and Dominik Bach for helpful comments on the manuscript.

## DATA AVAILABILTY

The data can be made available upon a reasonable request to the corresponding author.

## AUTHOR CONTRIBUTIONS

S.F. contributed to designing the experiments, data acquisition, data analysis, and writing the manuscript; J.D. contributed to data acquisition and data analysis; M.S. contributed to data analysis; and W.T. contributed to designing the experiment and writing the manuscript.

